# β1 integrin is a sensor of blood flow direction

**DOI:** 10.1101/511261

**Authors:** Ioannis Xanthis, Celine Souilhol, Jovana Serbanovic-Canic, Hannah Roddie, Antreas C. Kalli, Maria Fragiadaki, Raymond Wong, Dhruv R. Shah, Janet A. Askari, Lindsay Canham, Nasreen Akhtar, Shuang Feng, Victoria Ridger, Jonathan Waltho, Emmanuel Pinteaux, Martin J. Humphries, Matthew T. Bryan, Paul C. Evans

**Author notes:** Address for correspondence: Professor Paul C Evans, Department of Infection, Immunity and Cardiovascular Disease, Medical School, University of Sheffield, Beech Hill Road, Sheffield S10 2RX, UK.

## Abstract

The ability of endothelial cells (EC) to sense blood flow direction is a critical determinant of vascular health and disease. Unidirectional flow induces EC alignment and vascular homeostasis, whereas bidirectional flow has pathophysiological effects. EC express several mechanoreceptors that can respond to fluid flow (shear stress) but the mechanism for sensing the direction of shearing force is poorly understood. We observed using *in vitro* flow systems and magnetic tweezers that β1 integrin is a key sensor of force direction because it is activated by unidirectional but not bidirectional shearing forces. Consistently, β1 integrin was essential for Ca^2+^ signalling and cell alignment in response to unidirectional but not bidirectional shear stress. β1 integrin activation by unidirectional force was amplified in EC that were pre-sheared in the same direction, indicating that alignment and β1 integrin activity has a feedforward interaction which is a hallmark of system stability. *En face* staining and EC-specific genetic deletion studies of the murine aorta revealed that β1 integrin is activated and is essential for EC alignment at sites of unidirectional flow but is not activated at sites of bidirectional flow. In summary, β1 integrin sensing of unidirectional force is a key mechanism for decoding blood flow mechanics to promote vascular homeostasis.

**SUMMARY STATEMENT:** We demonstrate using flow systems, magnetic tweezers and knockout mice that β1 integrin is a sensor of shear stress direction in endothelial cells. β1 integrin is activated by unidirectional hemodynamic shear force leading to calcium signalling and cell alignment, but it is not activated by bidirectional force.

## INTRODUCTION

Although multiple mechanoreceptors have been identified, the fundamental mechanisms that cells use to sense the *direction* of force remain largely unknown. Arteries are exposed to mechanical forces of differing direction and magnitude via the action of flowing blood which generates shear stress (mechanical drag) on endothelial cells (EC) which line the inner surface. Notably, atherosclerosis, the major cause of mortality in Western societies, develops at branches and bends of arteries that are exposed to disturbed non-uniform flow (Kwak et al., 2014). These flow fields are remarkably complex and include flow that oscillates in direction (bidirectional), secondary flows that are perpendicular to the main flow direction, and low velocity flows. By contrast, artery regions that are exposed to non-disturbed unidirectional shear stress are protected. The direction of shear stress has profound effects on EC physiology. Unidirectional shear stress induces EC alignment accompanied by quiescence, whereas bidirectional and other non-uniform shear stress profiles do not support alignment (Ajami et al., 2017; Feaver et al., 2013; Sorescu et al., 2004; Wang et al., 2013; Wang et al., 2006; Wu et al., 2011). EC express several mechanoreceptors including integrins, ion channels, the glycocalyx, primary cilia and G-protein coupled receptors (Baeyens et al., 2014; Chen et al., 2015; Chen et al., 1999; Friedland et al., 2009; Givens and Tzima, 2016; Matthews et al., 2006; Shyy and Chien, 2002; Tzima et al., 2001; Tzima et al., 2005). However, the mechanisms that allow cells to decode the direction of shear stress are poorly characterised and a key question in vascular biology.

The integrin family of α/β heterodimeric adhesion receptors mediate adhesion of cells to neighbouring cells or to extracellular matrix via interaction with specific ligands. This process involves quaternary structural changes in integrin heterodimers, whereby a low affinity, bent configuration is converted to a high affinity, extended form (Friedland et al., 2009; Li et al., 2017; Puklin-Faucher et al., 2006; Puklin-Faucher and Sheetz, 2009). The ability of integrins to sense and respond to force is essential for cell shape, tissue architecture, cell migration and other fundamental processes. In the vasculature, the influence of flow on EC physiology involves shear stress-mediated activation of integrins (Tzima et al., 2001), which engage with extracellular matrix thereby triggering outside-in signalling (Bhullar et al., 1998; Chen et al., 2015; Chen et al., 1999; Jalali et al., 2001; Orr et al., 2006; Orr et al., 2005; Shyy and Chien, 2002; Tzima et al., 2001). Activation of α5β1 integrins by shear stress leads to Ca^2+^ signalling (Buschmann et al., 2010; Loufrani et al., 2008; Matthews et al., 2006; Thodeti et al., 2009; Yang et al., 2011), which in turn promotes vascular quiescence and health (Urbich et al., 2002) (Urbich et al., 2000) (Luu et al., 2013). One model for the role of integrins in shear stress signalling is that tension generated at the apical surface is transmitted through the cytoskeleton to integrins localized to the basal surface, thereby inducing structural changes that enhances their affinity for extracellular matrix ligands (Bhullar et al., 1998) (Orr et al., 2006; Orr et al., 2005) (Puklin-Faucher and Sheetz, 2009) (Tzima et al., 2001). However, studies using magnetic beads applied to the surface of cultured EC indicate that the apical pool of integrin can also respond to mechanical force (Matthews et al., 2006), a finding that we have explored further in this manuscript.

Here we studied the fundamental mechanism used by EC to sense the direction of mechanical force. This topic has translational significance because the mechanoreceptors that sense unidirectional protective force could be potentially targeted therapeutically to treat atherosclerosis. Although there is abundant evidence for the role of β1 integrin in mechanotransduction, its potential role in sensing the direction of flow has not been studied previously. We concluded that β1 integrins are essential for EC sensing of force direction since they activated by unidirectional force to drive EC alignment but were not activated by bidirectional force. Thus β1 integrins are the first example of a receptor that is activated specifically by unidirectional flow.

## RESULTS

### β1 integrin is activated by unidirectional but not by bidirectional shearing force

To investigate whether β1 integrin responds to a specific flow direction, we exposed cultured human umbilical vein EC (HUVEC) to shear stress (15 dyn/cm^2^) that was either unidirectional or bidirectional (1 Hz). Staining with the 12G10 antibody, which specifically binds to the high affinity, extended β1 conformer, revealed that β1 integrins were activated by exposure to unidirectional but not bidirectional flow (Fig. 1A). Since flow can alter the transport of materials as well as local mechanics, we used a magnetic tweezers platform to apply force directly to β1 integrins and determine whether force direction regulates their activity. An electromagnet was built in-house and coupled to a fluorescence microscopy platform fitted with an incubation chamber heated to 37°C, enabling live-cell imaging during operation of the tweezers. Passing current through copper coil pairs generated a magnetic field that was concentrated close to the sample by the corresponding pole piece. In this study, poles 1 and 2 were used to generate unidirectional or bidirectional forces (but poles 3 and 4 were not used; Fig. 1B). Computational modelling revealed that the force generated at the centre of the imaging area of the microscope was 16 pN, with a 7.5% (1 pN) variation across the imaging area. Forces along the y- and z-directions were negligible (< 0.1% of the total force), so the force generated by the tweezers was directed almost entirely along the x-direction, towards the activated pole (Fig. 1C, Fig. 1D). As the force was parallel to the stage, it mimicked the shearing action experienced by receptors under flow (Fig. 1E). The generation of force was validated by observing the movements of suspended paramagnetic beads (e.g. Supplemental Video 1).

**Figure 1.**
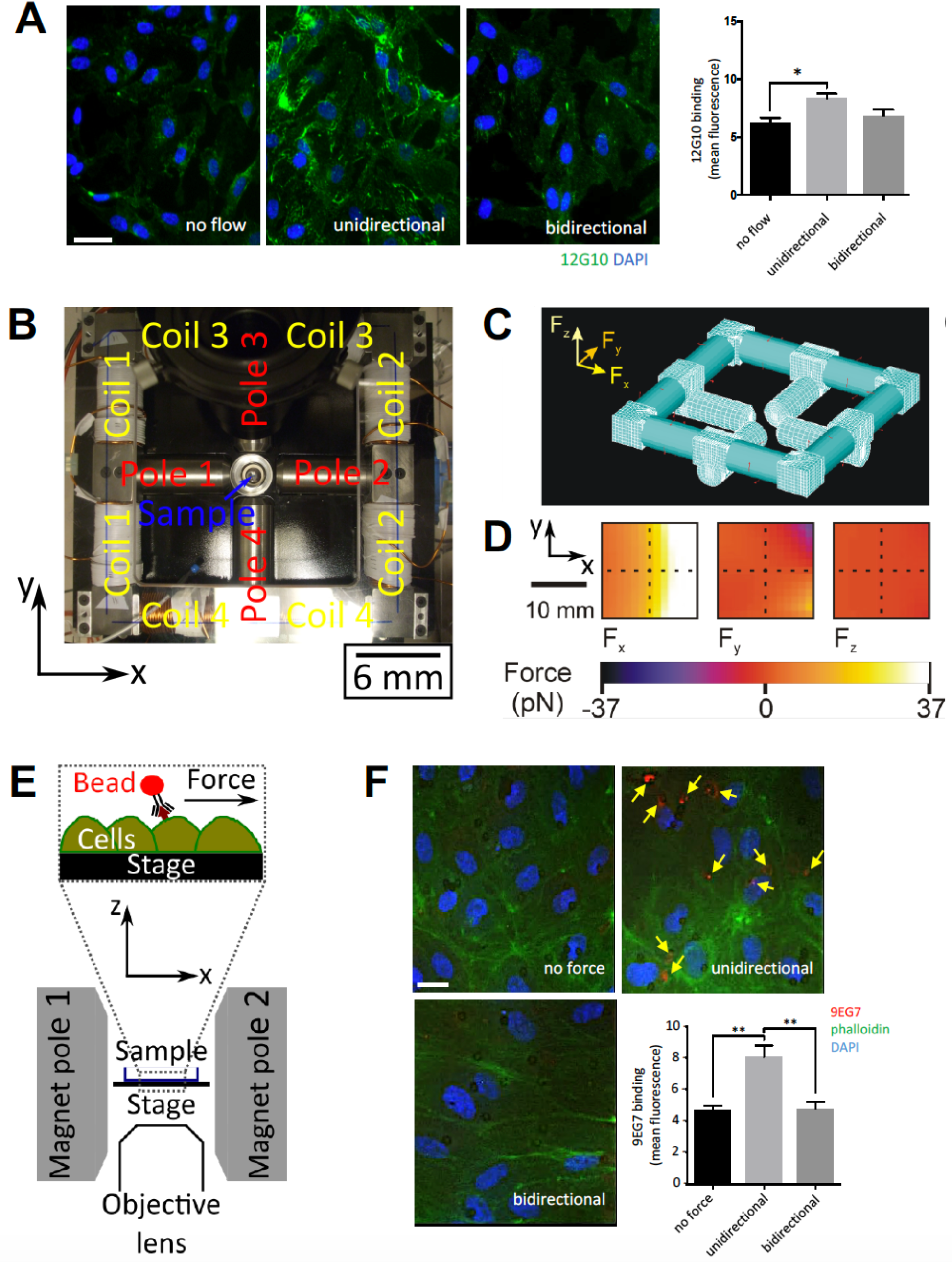
β1 integrins are activated by unidirectional but not bidirectional shearing force. (A) HUVECs were exposed to unidirectional or 1 Hz bidirectional flow for 3 min or remained under static conditions. Cells were stained with antibodies targeting active β1 integrins (12G10; green) and DAPI (nuclei; blue). Representative images and 12G10 mean fluorescence values ± SEM are shown. Scale bar: 10 μm. Data were pooled from 5 independent experiments. Differences were analysed using a one-way ANOVA with Tukey’s test for multiple comparisons. *p<0.05. (B) Photograph of the magnetic tweezers platform showing which coil pairs activate which pole, the sample position and the coordinate system. (C) The finite element mesh used in the ANSYS model of the electromagnet is shown. (D) The modelled forces generated along the x-, y- and z-directions (F_x_, F_y_, F_z_ respectively) within the plane of the stage position under when 10 A applied to coil 2 are shown. The hatched lines indicate the imaging position. (E) A cross-section schematic diagram of the sample position, highlighting the direction of the applied force when pole 2 is activated. (F) 4B4-coated magnetic beads (targeting the βI domain of inactive β1 integrin) were incubated with HUVEC prior to the application of unidirectional or 1 Hz bidirectional force (~16 pN) for 3 min. As a control, beads remained under no force. β1 integrin activation was quantified by immunostaining (9EG7; red) with co-staining of F-actin (phalloidin; green) and nuclei (DAPI; blue). Scale bar: 10 μm. Data were pooled from 4 independent experiments. Differences were analysed using a one-way ANOVA with Tukey’s test for multiple comparisons. **p<0.01.

The influence of force direction on β1 integrin activation was assessed by incubating HUVEC with superparamagnetic beads coated with antibodies that target the βI domain of the inactive β1 conformer (4B4 antibodies) prior to the application of unidirectional or bidirectional forces. After 3 min force, β1 integrin activation was quantified by staining using 9EG7 antibodies that specifically recognize the extended active conformer (Byron et al., 2009; Mould et al., 1995; Su et al., 2016). Since the shear stress generated by the bead is given by the force/contact area, we estimate the bead produces between 10-15 dyn/cm^2^, assuming that between 1/4 and 1/6 of the surface area of the bead is in contact with the cell. This is comparable magnitude to the shear stress in human arteries. It was observed that the application of unidirectional force enhanced 9EG7 binding indicating that mechanical activation of β1 integrin is induced by unidirectional force, whereas bidirectional force had no effect (Fig. 1F). Therefore, unidirectional force converts β1 integrin into an extended active conformer whereas bidirectional force does not.

### The βI domain of β1 integrin senses unidirectional force

To determine the regions of β1 integrin that are responsible for force sensing, we applied force using monoclonal antibodies that target specific domains (Byron et al., 2009) (Fig. 2A). Activation of β1 integrin by mechanical force is known to induce Ca^2+^ signalling (Matthews et al., 2006) and we therefore used Ca^2+^ accumulation as a readout using the fluorescent Ca^2+^ reporter (Cal-520). Force did not cause detachment of beads from cells in these experiments (Supplementary Fig. 1). The application of unidirectional force to the βI domain of the inactive form (via mab13 or 4B4 antibodies) or to the active extended form (via TS2/16 or 12G10 antibodies) induced Ca^2+^ accumulation (Figs. 2B and 2C), whereas force application to the hybrid (HUTS4), PSI (8E3), EGF-like (9EG7) domains or membrane-proximal region (K20) had no effect (Fig. 2B). Of note, the βI domain discriminated between different patterns of force because it induced Ca^2+^ signalling in response to unidirectional but not bidirectional force (Fig. 2C). As a control, it was demonstrated that the application of force to poly-D lysine-coated beads, which bind negatively charged molecules, had no effect on Ca^2^ levels (data not shown). Thus, it was concluded that the βI domain of β1 integrin is a sensor of force direction; it responds specifically to unidirectional force leading to activation of β1 integrin and downstream Ca^2+^ signalling. Moreover, our observation that force can promote signalling when applied to pre-activated extended forms of β1 integrin implies that tension is transmitted through β1 integrin to the cell during signal transduction.

**Figure 2.**
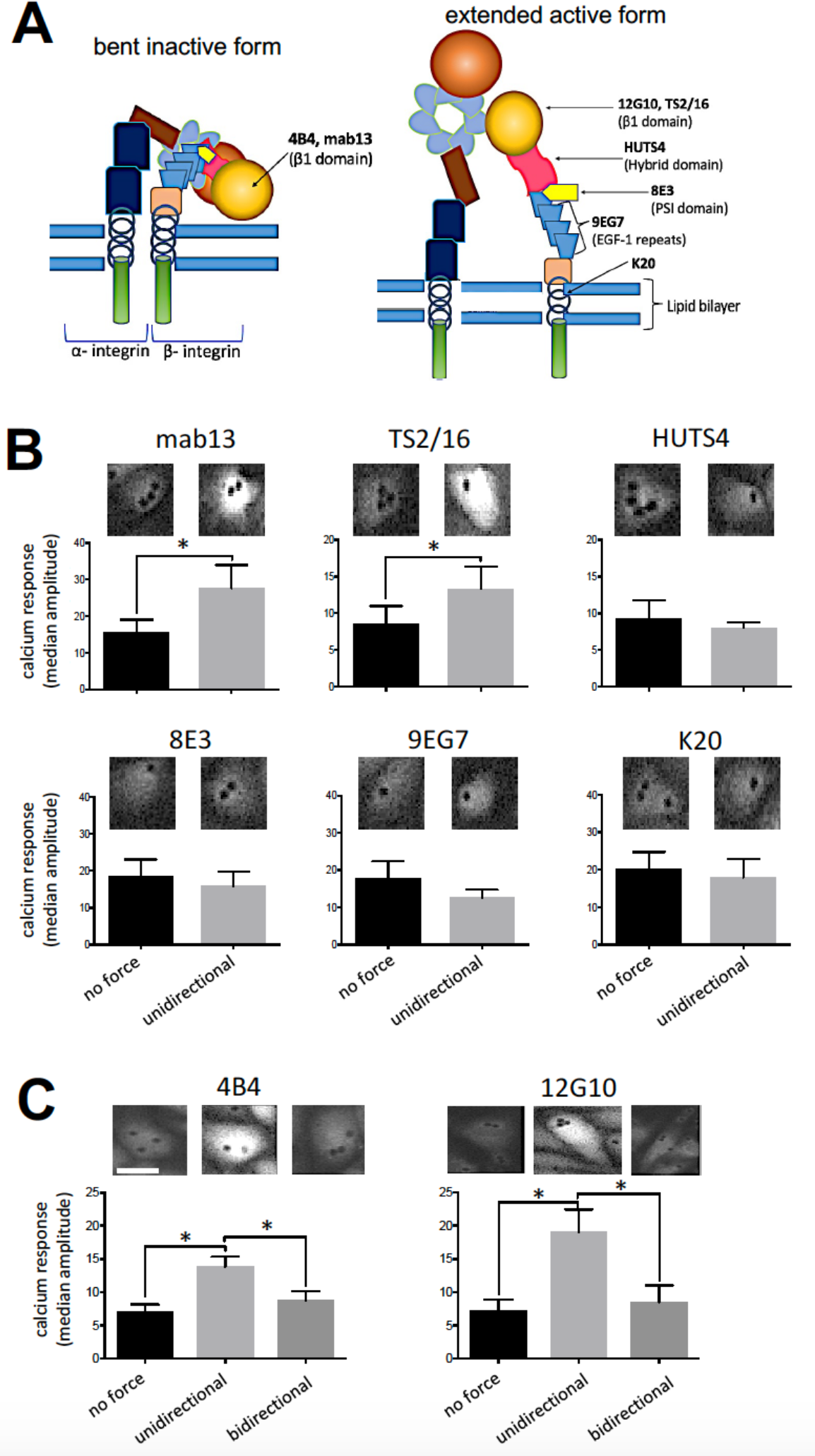
β1 integrins sense unidirectional force via the βI domain. (A) Schematic representation of domains and antibody binding sites on bent inactive or active β1 integrin. (B) HUVECs were loaded with Cal-520 and then incubated with beads coated with mab13, TS2/16, HUTS4, 8E3, 9EG7 or K20 antibodies. Beads were exposed to unidirectional force (~16 pN) or no force as a control. (C) HUVECs were loaded with Cal-520 and then incubated with beads coated with antibodies targeting inactive (4B4) or active (12G10) β1 integrins. Beads were exposed to unidirectional force (~16 pN), bidirectional force (1Hz ~16 pN) or no force. (B, C) Calcium responses were recorded for 3 min using fluorescence microscopy. Representative images are shown. Data were pooled from 5 independent experiments. The median amplitude of the first peak of the calcium response was calculated. Values are shown as means ± SEM and differences were tested using a two-tailed paired t-test (B) or one-way ANOVA test, with Tukey’s test for multiple comparisons (C). *p<0.05.

The mechanism of integrin activation by shear stress was also studied by steered molecular dynamic (SMD) simulations. The αVβ3 ectodomain structure (PBD:3IJE; (Xiong et al., 2009)) was used as a surrogate because a 3D structure of the β1 integrin ectodomain is not available. However, we can extrapolate these data to β1 integrin because there is high structural similarity between the β3 and β1 subunits and they have similar fold in the inactive state. Previous studies demonstrated that a pulling force applied tangentially to the membrane induces integrin activation by unfolding and increases the angle of the βI/hybrid domain hinge subsequently leading to αβ leg separation (Chen et al., 2012) (Puklin-Faucher et al., 2006). However, to mimic the effects of shear stress we applied force (200 kJ mol^-1^ nm^-1^) *parallel* to the membrane to the βA domain and demonstrated that it converted the bent inactive form into an extended form that was tilted relative to the membrane (Fig. 3 and Supplemental Video 2). Due to the size of this system (~1.5M atoms) we have used forces that are higher than physiological levels. For this reason, our simulations cannot provide information about the timescale of integrin activation by shear stress however they support our magnetic tweezers data showing that force applied parallel to the membrane can cause integrin activation.

**Figure 3.**
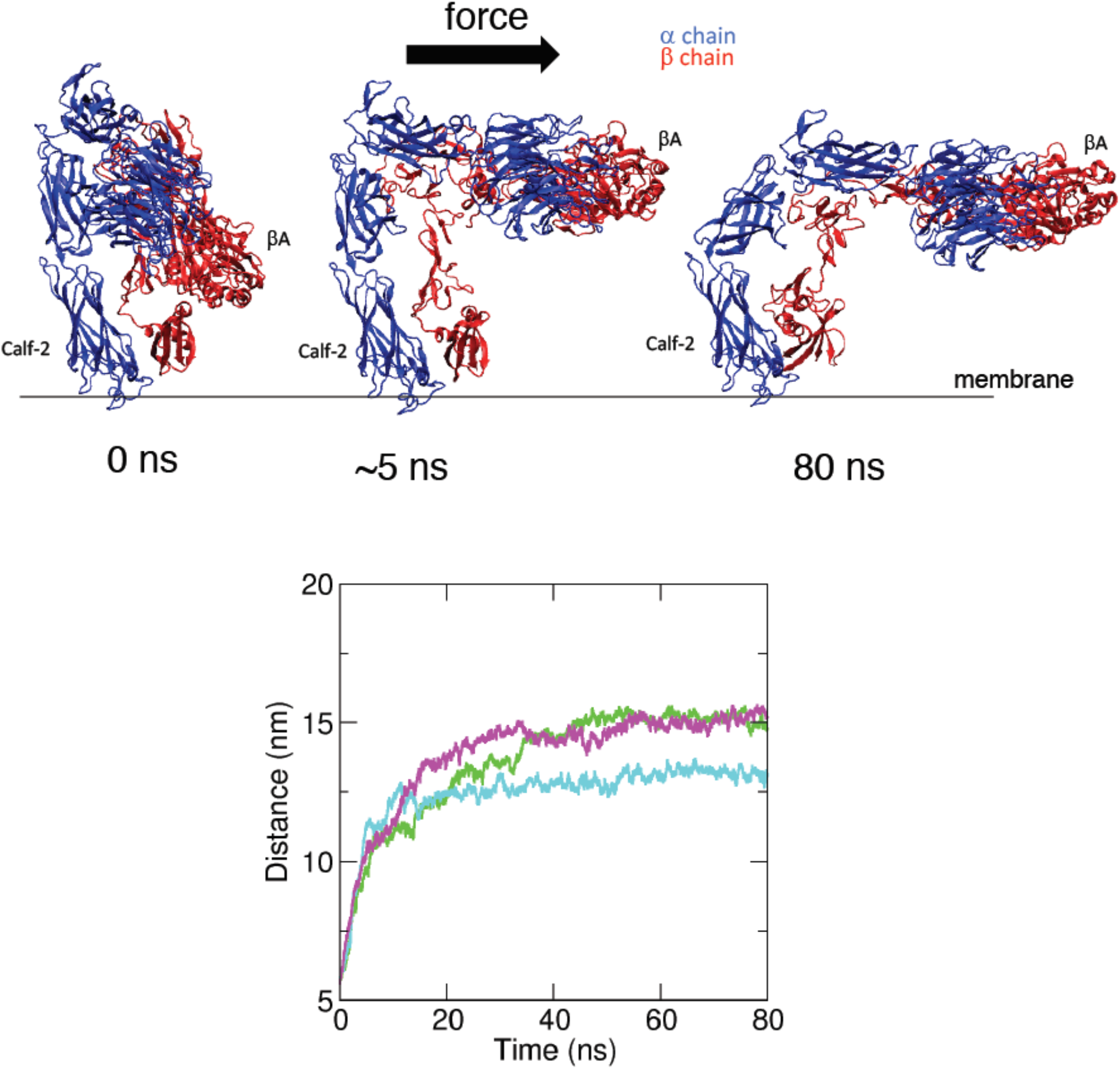
SMD simulations of integrin structural rearrangements in response to mechanical force. SMD simulations were performed using the αVβ3 integrin ectodomain. A force (200 kJ mol^-1^ nm^-1^) parallel to the membrane was applied on the βA domain of the head region of the inactive form. The constant application of force triggered intramolecular structural rearrangements and extension of the molecule in three independent simulations. The structure is shown at the beginning of the simulations and after the application of force for 5 ns or 80 ns. αV is shown in blue and β3 integrin is shown in red (upper panels). Structural rearrangements were quantified by measuring the distance between the Calf-2 and βA domains over the course of the simulation (lower panels). The coloured lines represent the three repeat simulations. Force consistently induced extension of the integrin heterodimer.

### β1 integrin elicits Ca^2+^ signalling in response to unidirectional flow but not bidirectional flow

Next, we investigated the mechanism by which β1 integrin converts unidirectional flow into Ca^2+^ signalling as this pathway is a pivotal regulator of EC physiology (Ando and Yamamoto, 2013). Imaging of HUVEC loaded with Cal-520 revealed Ca^2+^ accumulation in the cytosol in response to unidirectional or bidirectional flow (Fig. 4A and Supplemental Video 3), indicating that both of these flow patterns drive Ca^2+^ signalling. However, β1 integrin blocking antibodies (P5D2) reduced Ca^2+^ accumulation in response to unidirectional flow but not bidirectional flow or static conditions (Fig. 4B). Thus, although both unidirectional and bidirectional flow activate Ca^2+^ signalling, β1 integrin is specifically required for the response to unidirectional flow. We next investigated whether β1 integrin-dependent Ca^2+^ signalling involves Piezo1 and TRPV4 since these Ca^2+^-permeable channels are known to sense shear stress (Kohler et al., 2006; Mendoza et al., 2010; Nguyen et al., 2014; Thodeti et al., 2009). HUvEcs were treated with specific siRNAs to silence Piezo1 or TRPV4 (Supplementary Fig. 2) prior to the application of unidirectional force via magnetic tweezers coupled to 12G10-coated superparamagnetic beads. Silencing of Piezo1 or TRPV4 significantly reduced Ca^2+^ accumulation indicating that both channels are involved in β1 integrin-dependent Ca^2+^ signalling (Fig. 4C). Consistently, β1 integrin-dependent Ca^2+^ signalling was significantly reduced by EGTA indicating a requirement for extracellular Ca^2+^ (Supplementary Fig. 3). By contrast, the response to bidirectional flow was only partially reduced by eGta and this difference was not statistically significant. We conclude that unidirectional force induces Ca^2+^ signalling via a mechanism that requires β1 integrin activation of Piezo1 and TRPV4 coupled to extracellular Ca^2+^, whereas bidirectional force signals via a β1 integrin-independent mechanism.

**Figure 4.**
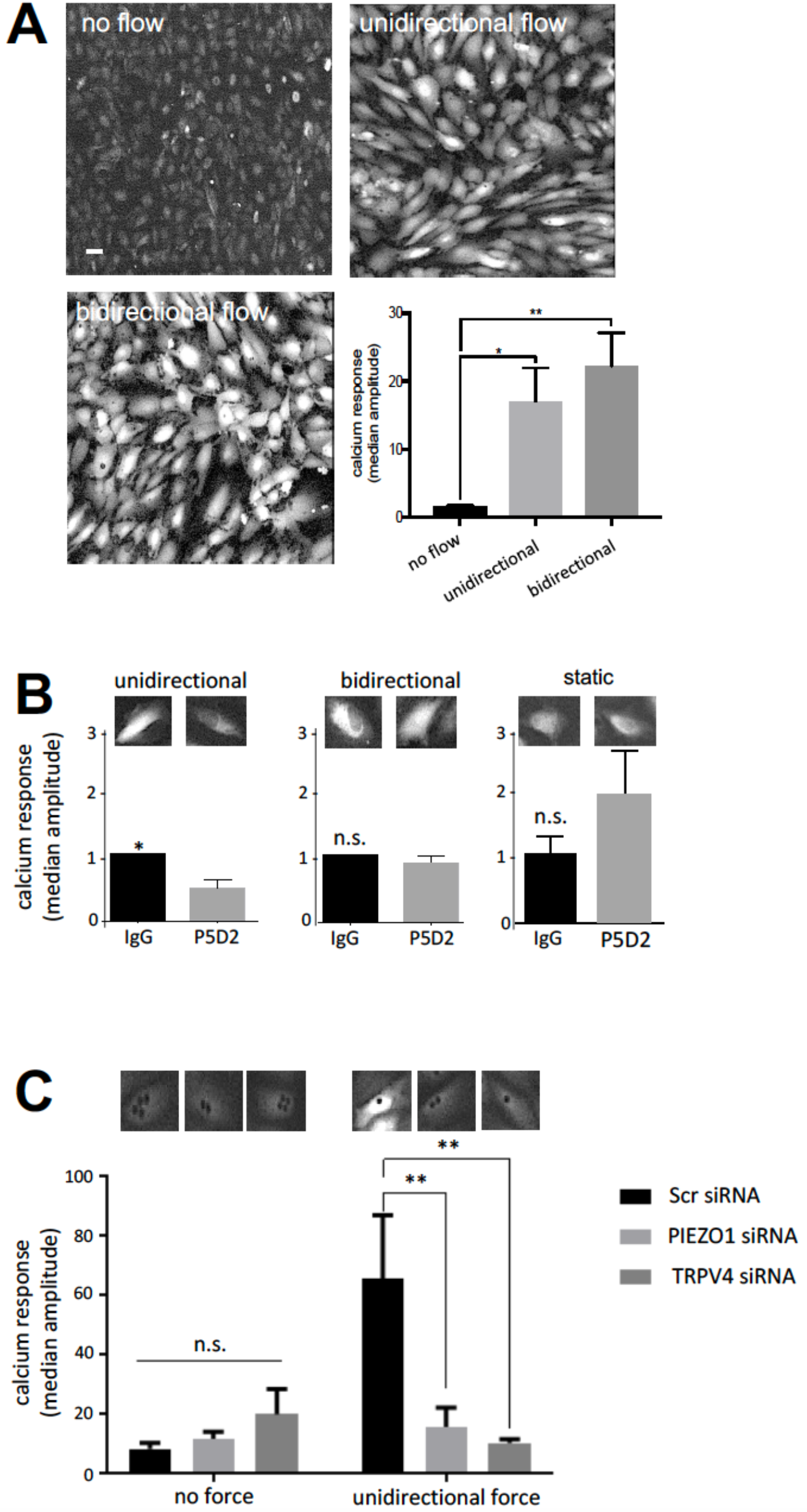
β1 integrins induce Ca^2+^ accumulation in response to unidirectional flow via Piezo1 and TRPV4. (A) HUVECs were loaded with the calcium fluorescent dye (Cal-520), then exposed to unidirectional or 1 Hz bidirectional flow and calcium responses were recorded for 3 min using fluorescence microscopy. Representative images are shown (scale bar: 10 μm). (B) HUVEC were loaded with Cal-520 and then incubated with P5D2 (β1 integrin blocking antibody) or with total isotype-matched mouse IgG as a control. They were then exposed to unidirectional or 1 Hz bidirectional flow or static conditions and calcium responses were recorded for 3 min. (C) HUVECs were transfected with siRNA targeting Piezo1, TRPV4 or with scrambled (Scr) sequences as a control. After 72 h, cells were loaded with Cal-520 and then incubated with beads coated with 12G10 antibodies targeting active β1 integrins. Beads were exposed to unidirectional force (~16 pN) or no force. Calcium responses were recorded for 3 min. Data were pooled from 5 (A), 6 (B) or 3 (C) independent experiments. The median amplitude of the first peak of the calcium response was calculated. Values are shown as means ± SEM and differences were tested using a one-way ANOVA test, with Tukey’s test for multiple comparisons (A), a two-tailed paired t-test (B) or a two-way ANOVA (C). *p<0.05, **p<0.01.

### Unidirectional shear stress induces a feedforward interaction between β1 integrin activation and cell alignment

Ca^2+^-signalling induces alignment of EC in the direction of flow which is essential for vascular homeostasis (Wang et al., 2013). To investigate the role of β1 integrins in this process we treated HUVEC with P5D2 activity blocking antibodies (or with non-binding antibodies as a control) during exposure to unidirectional or bidirectional flow. EC aligned specifically in response to unidirectional flow and this was blocked by P5D2 demonstrating an essential role for β1 integrin activation (Fig. 5A). Since EC polarity alters their response to flow (Wang et al., 2013), we investigated whether EC alignment can influence β1 integrin sensing of mechanical force. This was tested by exposing EC to shear stress and subsequently measuring the effects of applying force through β1 integrin either in the same direction as the flow, or in the opposite direction or perpendicular to the direction of flow. We observed an anisotropic response with faster signalling when force was applied in the same direction as the flow and slower responses when force was applied in the opposite direction or tangentially (Fig. 5B). Thus, unidirectional force sensing by β1 integrins is enhanced in cells that are aligned with flow, indicating a feedforward interaction between β1 integrin activation and cell alignment.

**Figure 5.**
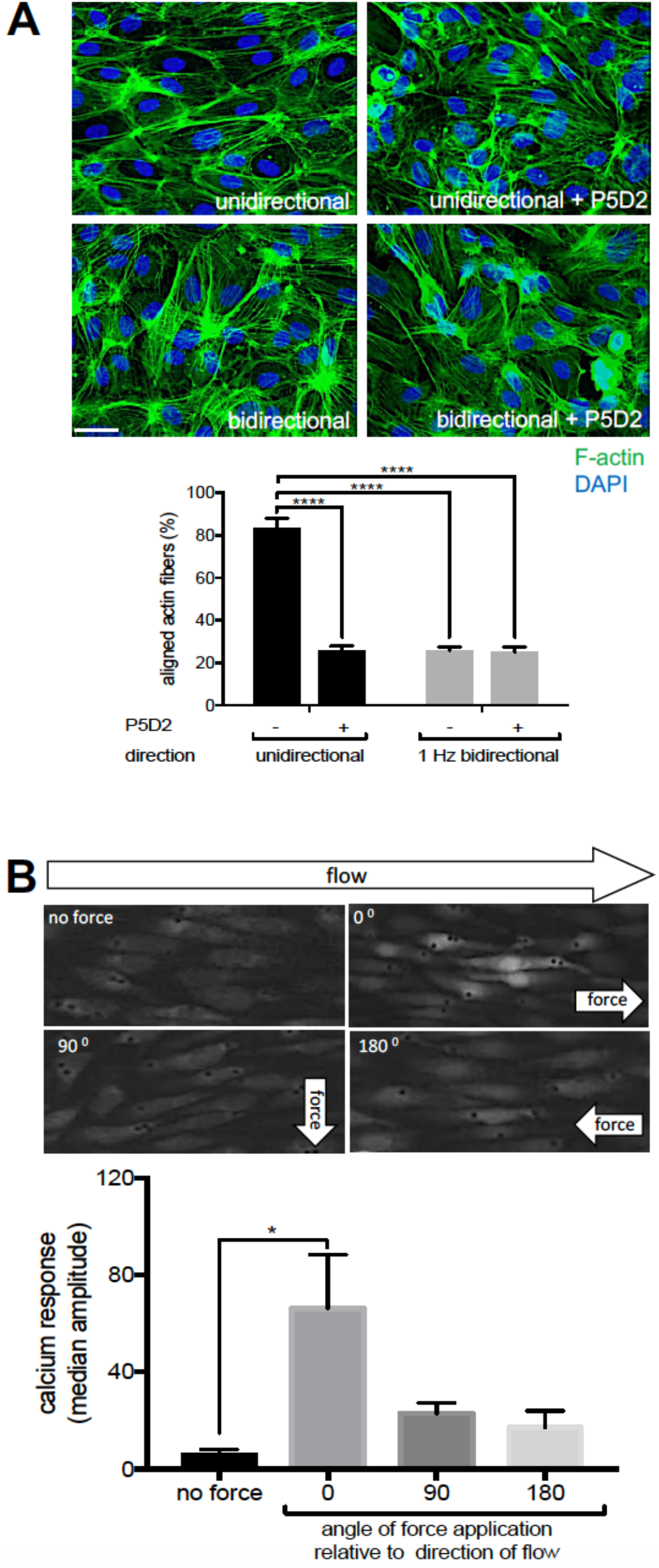
β1 integrin activation and cell alignment response have a feedforward interaction under unidirectional flow. (A) HUVECs pre-treated under static conditions with P5D2 (β1 integrin blocking antibody) or control IgG were exposed to unidirectional or 1 Hz bidirectional flow for 24 h and then stained with FITC-phalloidin (green, actin fibres) or DAPI (blue, nuclei). The proportion of cells with actin fibres aligned with the major cell axis (within 30°) was calculated. Scale bar: 10 μm. Mean values ± SEM are shown. (B) HUVEC were exposed to unidirectional flow for 72 h and then maintained under static conditions. They were loaded with the calcium fluorescent dye Cal-520 and incubated with beads coated with 12G10 antibody. Unidirectional force was then applied either in the same direction as the pre-shearing (0°), or tangentially (90°) or in the opposite direction (180°). Calcium responses were recorded for 3 min using fluorescence microscopy. Representative images are shown. Data were pooled from 3 independent experiments. The median amplitude of the first peak of the calcium response was calculated. Differences were analysed using a 2-way ANOVA test (A) or a one-way ANOVA test with Tukey’s test for multiple (B). *p<0.05, ****p<0.0001.

### β1 integrin is essential for EC alignment at sites of unidirectional shear *in vivo*

To assess whether β1 integrin activation correlates with flow direction *in vivo*, we quantified it at the outer curvature of the murine aortic arch which is exposed to unidirectional flow and at the inner curvature which is exposed to disturbed bidirectional flow (Suo et al., 2007). *En face* staining using the 9EG7 antibody followed by super-resolution confocal microscopy revealed that activation of β1 integrin was significantly higher at the outer curvature of the aortic arch compared to the inner curvature (Fig. 6A and Supplemental Video 4). A portion of active β1 integrin at the outer curvature was observed at the apical surface which is in direct contact with flowing blood (Fig. 6A). Our observation that β1 integrin is activated preferentially at a site of unidirectional flow in the murine aorta is consistent with the finding that it is activated exclusively by unidirectional shear stress in cultured EC (Figs 1 and 2).

**Figure 6.**
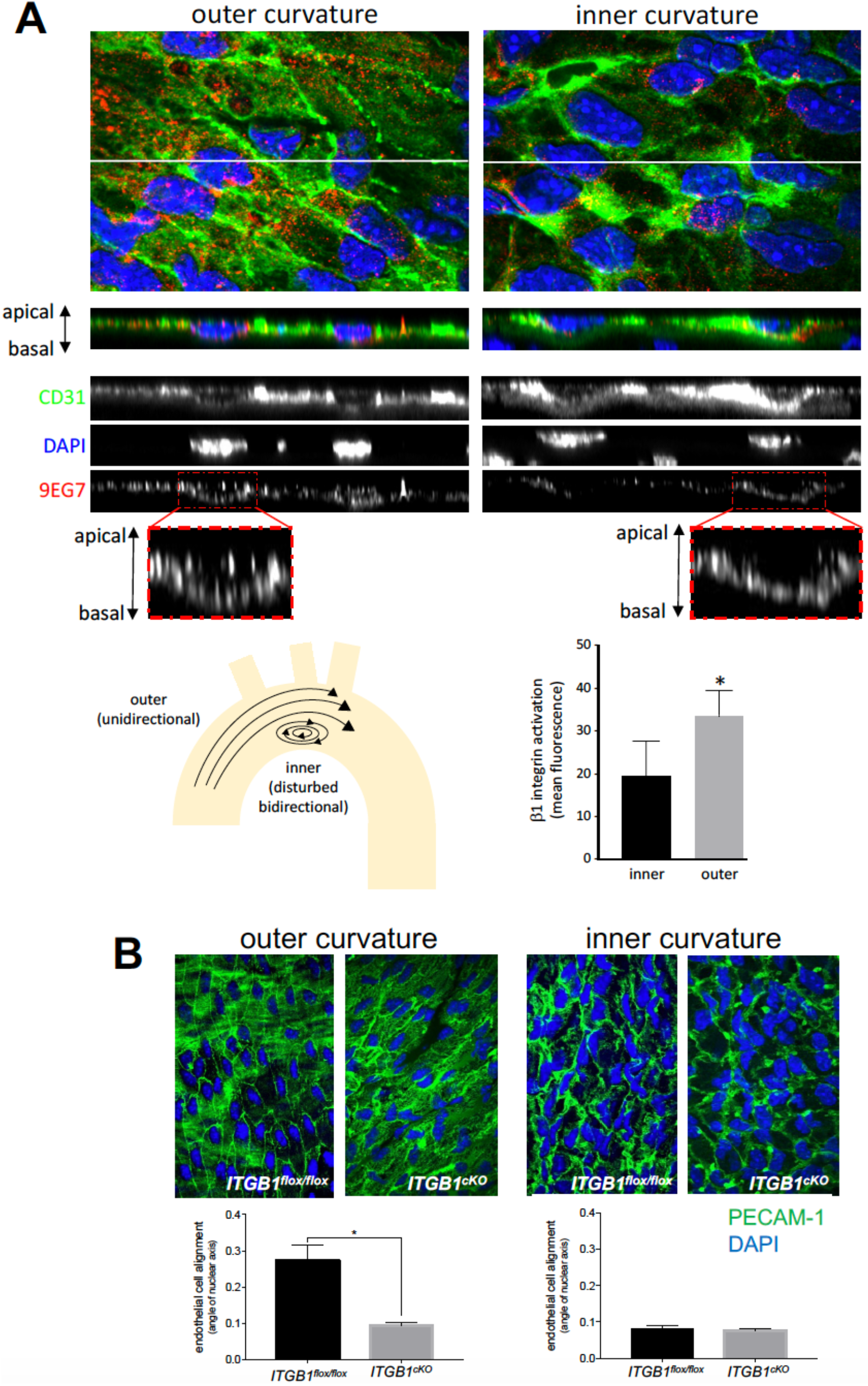
β1 integrin activation is essential for EC alignment at sites of unidirectional flow *in vivo*. (A) Mouse aortic arches were stained *en face* with antibodies targeting active β1 integrins (9EG7; red). EC were co-stained using anti-PECAM-1 antibodies (green) and nuclei were counterstained using DAPI (blue). Fluorescence was measured at the outer curvature (unidirectional flow) and inner curvature (bidirectional flow) regions using super-resolution confocal microscopy. Representative z-series stacks of images are shown (a, apical surface; b, basal surface). Note that 9EG7 stained apical and basal surfaces at the outer curvature but was restricted to the basal side at the inner curvature. Levels of active β1 integrins at the apical and basal surfaces were calculated by quantifying the ratio of fluorescence from 9EG7 (active form) and Mab 1997 (total β1 integrin). (B) Ec alignment was quantified at the outer curvature (unidirectional flow) and inner curvature (disturbed bidirectional flow) of the aortic arch by *en face* staining of PECAM-1 (green) and DAPI (nuclei; blue) in *itgb1*^fl/fl^ or *itgb1*^cKO^ mice (n=3). EC alignment was quantified by measuring the angle of the major axis of the nucleus. Mean fluorescence values ± SEM are shown (n=3 mice). Differences were analysed using either a two-tailed paired t-test (A) or a two-tailed, paired t-test (B). *p<0.05, ****p<0.0001.

To study the function of β1 integrin *in vivo* we deleted it conditionally from adult EC noting that deletion from embryonic EC is lethal (Carlson et al., 2008; Lei et al., 2008; Tanjore et al., 2008; Zovein et al., 2010). Conditional genetic deletion of β1 integrin from EC was confirmed by *en face* staining using anti-β1 integrin antibodies (Supplementary Fig. 4). Deletion of β1 integrin resulted in significantly reduced EC alignment at the outer curvature (unidirectional flow) but did not alter EC morphology at the inner curvature of the murine aortic arch (disturbed bidirectional flow) which were non-aligned in both wild-type and β1 integrin knockout mice (Fig. 6B). It should be noted that the β1 integrin knockout does not discriminate between apical and basal pools of integrin, however it can be used to support the concept that β1 integrin responds specifically to unidirectional flow. Collectively, our data demonstrate that β1 integrin activation by unidirectional shear stress is an essential driver of EC alignment.

## DISCUSSION

### Endothelial sensing of flow direction: role of β1 integrins

The ability of EC to sense the direction of blood flow is essential for vascular health and disease (Wang et al., 2013). It underlies the focal distribution of atherosclerotic lesions which develops at parts of arteries that are exposed to complex flow patterns including bidirectional flow but does not develop at sites of unidirectional flow. It is well established that EC sense the shearing force generated by flow via multiple mechanoreceptors including the VE-cadherin/PECAM-1/VEGFR2 trimolecular complex (Tzima et al., 2005), Piezo1 (Li et al., 2014) and several others. However, the molecular mechanisms that convert directional cues into specific downstream responses are poorly understood. Recent studies indicate that PECAM-1 can sense both unidirectional and disturbed flow leading to the transmission of protective and inflammatory signals accordingly. Thus PECAM-1 knockouts have a fascinating phenotype characterised by enhanced lesions at sites of unidirectional flow and reduced lesion formation at sites of disturbed flow (Goel et al., 2008; Harry et al., 2008). On the other hand, the transmembrane heparan sulphate proteoglycan syndecan-4 is required for EC alignment under shear stress but is dispensable for other mechanoresponses, indicating a role in sensing of flow direction (Baeyens et al., 2014).

Here we conclude that β1 integrins are sensors of force direction through the following lines of evidence: (1) β1 integrin converts from a bent inactive form to an extended active conformer in response to unidirectional but not bidirectional shearing force, (2) SMD simulations revealed that force applied parallel to the membrane can cause structural rearrangements leading to β1 integrin extension, (3) unidirectional shearing force induces Ca^2+^ signalling via a β1 integrin-dependent mechanism whereas the response to bidirectional force is independent from β1 integrin, (4) silencing of β1 integrin prevented alignment of cultured EC exposed to unidirectional shear stress but did not alter the morphology of cells exposed to bidirectional shear, (5) β1 integrin was activated specifically at sites of unidirectional shear stress in the murine aorta, (6) deletion of β1 integrin from EC reduced EC alignment at sites of unidirectional shear stress in the murine aorta but didn’t alter morphology at sites of disturbed flow.

### Is β1 integrin a direct sensor of flow?

There is abundant evidence that integrins can respond to flow *indirectly* via signals elicited from mechanoreceptors including PECAM-1 (Collins et al., 2012) and Piezo1 (Albarran-Juarez et al., 2018). Thus, flow causes activation of integrins on the basal surface of EC which subsequently engage with ligand and trigger outside-in signalling (Orr et al., 2006; Orr et al., 2005; Tzima et al., 2001). However, our observations suggest that an apical pool of β1 integrin exists that is activated by unidirectional shear stress to induce downstream signalling and cell alignment. Our data are consistent with a previous study in which mechanical signalling of apical integrins was induced with magnetic tweezers (Matthews et al., 2006). They also resonate with biochemical and electron microscopy studies that detected β1 integrin and other integrins at the apical surface of EC (Conforti et al., 1992; Conforti et al., 1991). However, they contrast with other studies that detected basal but not apical pools of β1 integrin using confocal microscopy (Li et al., 1997; Tzima et al., 2001). The reason for this discrepancy is uncertain, but may relate to our use of super resolution microscopy, which can delineate apical and basal surfaces of EC (<1 μm depth) more accurately than conventional confocal microscopy techniques. It is important to note that our observation that apical β1 integrin can respond to force does not preclude the important and well established role for basally-located integrins and we suggest that both pools contribute to flow sensing. Indeed, it is plausible that the function of α5β1 heterodimers varies according to their localisation on basal or apical surfaces since Albarrán-Juárez et al found that basally-located integrin is activated in response to disturbed flow (Albarran-Juarez et al., 2018), whereas we found that apically-located integrin is activated exclusively by unidirectional flow. The mechanisms that EC use to integrate these divergent downstream signals from apical and basal pools of β1 integrin should now be investigated further.

### Mechanism of unidirectional flow sensing and cell alignment

Using magnetic tweezers, we determined that unidirectional shearing force induces downstream signals via a 2-stage process. Firstly, it converts the bent inactive form of β1 integrin into an extended form. Secondly, the extended form of β1 integrin transmits force to the cell to elicit downstream signalling. At the basal surface of cells, β1 integrin is anchored to extracellular matrix and therefore can transmit tension to the cell (Friedland et al., 2009; Nordenfelt et al., 2016; Zhu et al., 2008). Although apical β1 integrins are not anchored to extracellular matrix, we hypothesize that they may function as a ‘sea anchor’ in cells exposed to flow, thereby allowing force to be transmitted to the cell. Thus, we propose that unidirectional shear stress induces tension in β1 integrin leading to downstream signalling whereas bidirectional shear stress is insufficient because it switches direction before tension can be established (Fig. 7). Our model is consistent with other studies that demonstrated that mechanical forces can activate β1 integrins independently from ligand binding (Ferraris et al., 2014; Petridou and Skourides, 2016).

**Figure 7.**
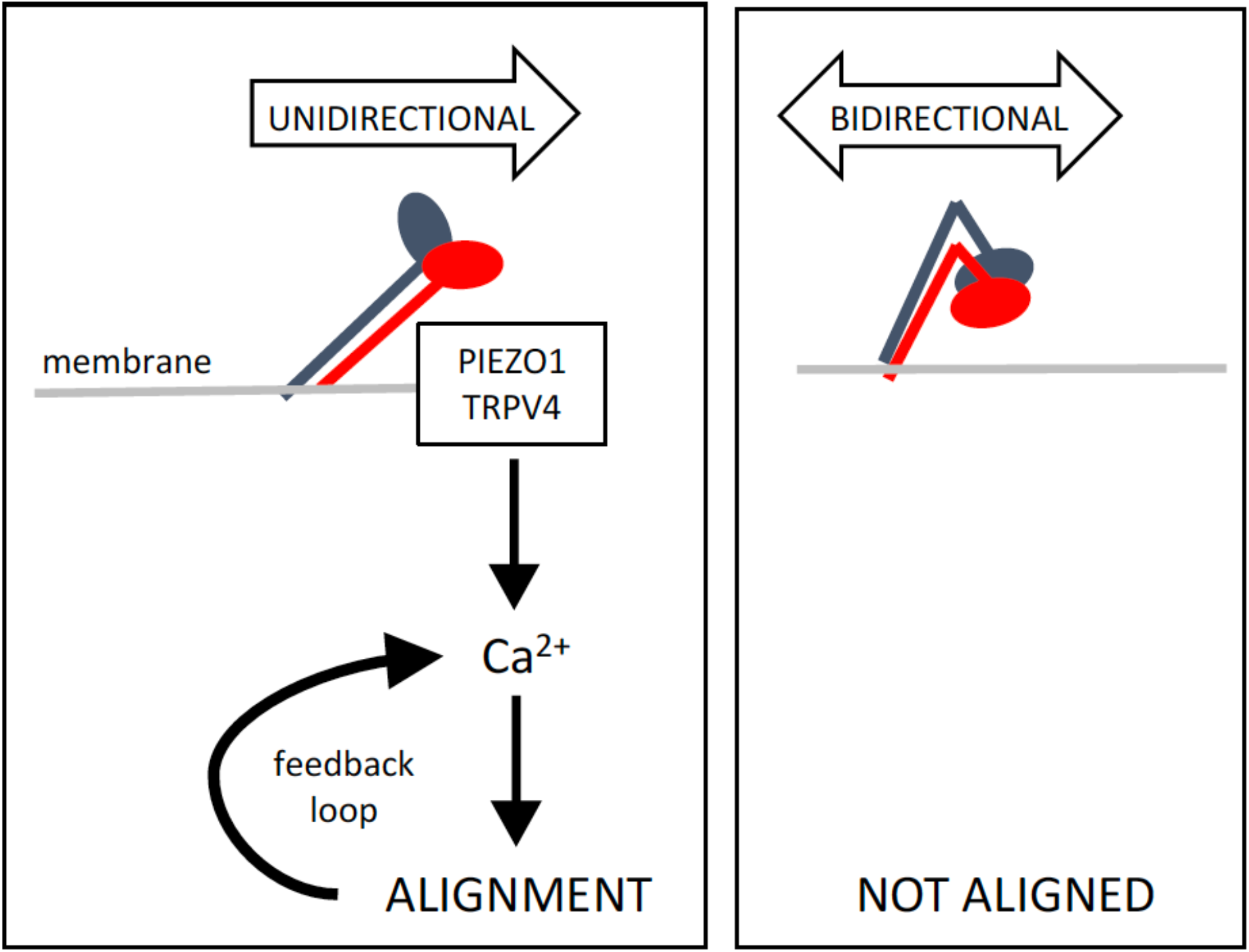
Model to explain the mechanism of β1 integrin sensing of flow direction. Unidirectional flow (left) induces structural changes in the ectodomain of β1 integrins causing them to extend. The integrin subsequently acts as a sea anchor thereby inducing the accumulation of Ca^2+^ via Piezo1 and TRPV4 leading to cell alignment. This forms a feedforward loop to enhance Ca^2+^ signalling therefore promoting physiological stability. Under bidirectional flow (right) β1 integrin is not activated.

We observed that β1 integrin sensing of unidirectional force induced Ca^2+^ accumulation via Piezo1 and TRPV4. These data are consistent with the known roles of Piezo1 (Li et al., 2014) and TRPV4 (Kohler et al., 2006) in shear sensing and with a previous report of crosstalk between β1 integrin and TRPV4 in sheared endothelium (Matthews et al., 2010). Our study also revealed that β1 integrins are essential for alignment of EC under unidirectional shear stress. Since Piezo1 and Ca^2+^ positively regulate EC alignment (Li et al., 2014) (Miyazaki et al., 2007) we propose that unidirectional force induces EC alignment via β1 integrin/ Piezo1/TRPV4-dependent Ca^2+^ signalling. The mechanisms of cross-talk between β1 integrins, Piezo1 and TRPV4 during endothelial responses to mechanical force should be studied further.

Interestingly, we observed that direction-specific β1 integrin signalling is anisotropic because it is enhanced in cells that are pre-aligned in the direction of force application but reduced in cells exposed to flow in the opposite direction or tangentially. The mechanism of anisotropy is uncertain but could involve flow-mediated alteration of actin dynamics which can modulate integrin orientation (Nordenfelt et al., 2017). Since β1 integrin drives EC alignment and *vice versa*, we conclude that a feedforward loop exists between β1 integrin activation and alignment. Feedforward systems are intrinsic to physiological stability and therefore the positive interaction between EC alignment and β1 mechanosensing is predicted to maintain long term vascular homeostasis at sites of unidirectional flow.

### Significance

Our study provides insight into the mechanisms that EC use to decode complex mechanical environments to produce appropriate physiological responses. Focussing on β1 integrin, we found that they are specific sensors of unidirectional flow driving downstream signalling and EC alignment. These findings suggest the exciting possibility that specific mechanical force profiles are sensed by specific mechanically-cognate receptors to elicit distinct downstream responses. Future studies should now identify the mechanoreceptors that sense other mechanical forces profiles e.g. bidirectional force. Our observation that EC responses to distinct force profiles can be modified by targeting specific mechanoreceptors has implications for the treatment of atherosclerosis which develops and progresses at sites of disturbed flow (Kwak et al., 2014).

## MATERIALS AND METHODS

### Antibodies

The following monoclonal antibodies that recognise β1 integrin were used: 12G10 (Abcam ab30394; specifically recognises the active form), 9EG7 (BD Pharmingen 553715; specifically recognises the active form), P5D2 (Abcam ab24693; blocks activation), 4B4 (Beckman Coulter 41116015), TS2/16, HUTS4, 8E3, K20, mab13 (Byron et al., 2009; Mould et al., 1995), mab1997 (Merck MAB1997). Rabbit anti-integrin β1 antibodies (ab179471), and antibodies recognising murine CD31 (Biolegend) and human CD144 (BD Pharmingen) were used.

### EC culture and application of shear stress

HUVECs were isolated using collagenase digestion and maintained in M199 growth medium supplemented with foetal bovine serum (20%), L-glutamine (4 mmol L^-1^), endothelial cell growth supplement (30 μg ml^-1^), penicillin (100 U μl^-1^), streptomycin (100 μg ml^-1^) and heparin (10 U ml^-1^). HUVECs (25×10^4^) were seeded onto 0.4 mm microslides (Luer ibiTreat, ibidi™) pre-coated with 1% fibronectin (Sigma) and used when they were fully confluent. Chamber slides were placed on the stage of an inverted light microscope (Nikon^®^ ECLIPSE Ti) enclosed in a Perspex box pre-warmed to 37°C. Unidirectional or 1 Hz bidirectional flow of 15 dynes cm^-2^ for the indicated time was applied using the ibidi™ syringe pump system. Pharmacological inhibition of β1 integrin activation was performed using 1-10 μg ml^-1^ P5D2 antibody (Abcam).

### Gene silencing and quantitative RT-PCR

Gene silencing was performed using small interfering (si) RNA sequences from Dharmacon (Piezo1 (L-020870-03), TRPV4 (L-004195-00)). A non-targeting control siRNA (D-001810-10) was used as a control. HUVECs were transfected using Neon Transfection System (Invitrogen) and following manufacturer’s instructions. Final siRNA concentration was 50 nM. To determine the efficiency of the knockdown, total RNA was extracted using RNeasy Mini Kit (QIAGEN) according to manufacturer’s protocol and 500 ng of total RNA was subjected to cDNA synthesis using iScript reverse transcriptase (Bio-Rad). Resulting cDNA was used as a template for quantitative RT-PCR (qRT-PCR) using gene-specific primers and SsoAdvanced Universal SYBR Green Supermix from Bio-Rad. Amplification of housekeeping gene HPRT was used as an internal control. Following primer sequences were used: HPRT (Forward 5’-TTG GTC AGG CAG TAT AAT CC-3’; Reverse 5’-GGG CAT ATC CTA CAA CAA AC-3’); Piezo1 (Forward 5’-GCC GTC ACT GAG AGG ATG TT-3’; Reverse 5’-ACA GGG CGA AGT AGA TGC AC-3’); TRPV4 (Forward 5’-CTA CGG CAC CTA TCG TCA CC-3’; Reverse 5’-CTG CGG CTG CTT CTC TAT GA-3’).

### Immunofluorescent staining of cultured EC

Activation of β1 integrin was assessed by immunofluorescent staining using 9EG7 or 12G10 antibodies and AlexaFluor488-conjugated secondary antibodies (Invitrogen). Imaging was performed using a fluorescence microscope (Olympus) or a super-resolution confocal microscope (Zeiss LSM880 AiryScan Confocal).

### Quantification of calcium responses

HUVECs seeded onto 1% fibronectin-coated 0.4 microslides (Luer ibiTreat, ibidi™) or 35 mm microdishes (μ-Dish 35 mm, ibidi™) were incubated with 50 μg Cal-520™ (AAT Bioquest^®^) and Pluronic^®^ F-127 (Invitrogen). For testing the effect of the directionality of force on pre-aligned cells, HUVECs were seeded onto 6-well plates with circular glass coverslips (13 mm diameter) attached to the periphery of the wells. The cells were exposed to flow for 72 hours using the orbital shaker model (Warboys et al., 2014) and the coverslips subsequently removed, placed into ibidi 35 mm microdishes and incubated with Cal520™ and Pluronic^®^ F-127 as described above. After incubation, cells were washed twice with sodium-buffered saline Ca^2+^ - containing media (134.3 mM NaCl, 5 mM KCl, 1.2 mM MgCl_2_, 1.5 mM CaCl_2_, 10 mM HEPES, 8 mM Glucose, pH 7.4) and maintained in this medium for subsequent experiments. Medium that lacked CaCl_2_ and included 0.4 mM EGTA was used for experiments requiring depletion of extracellular Ca^2+^. To measure calcium responses, Cal-520™ fluorescence was recorded using an inverted fluorescence microscope (Nikon Eclipse *Ti*) coupled to a photometrics CoolSnap MYO camera (180 consecutive images of the cells were recorded, with each image to be taken every second). Mean fluorescence values were extracted for single cells using ImageJ software (1.48v) and plotted against time to generate a kinetic profile. The amplitude of the first peak was calculated by deducting the minimum intensity value from the maximum intensity value and then dividing by the minimum intensity value (see Supplementary Fig. 5 for examples). Data were pooled from > 50 cells to generate a median peak amplitude.

### Coating of magnetic beads

Superparamagnetic beads (4.5 μm diameter; 1×10^7^; Dynabeads) conjugated to goat antimouse IgG, sheep anti-rat IgG antibodies (Invitrogen; 200 μg ml^-1^) were coated non-covalently with the antibodies of interest or covalently to Poly-D lysine (Sigma; 200 μg ml^-1^). They were washed with phosphate buffered saline containing 0.1% (w/v) bovine serum albumin and 0.5 M EDTA (pH 7.4) and resuspended in serum-free M199 media.

### Magnetic tweezers

A mild steel-cored electromagnet was built in-house and set into the stage of an inverted fluorescence microscope (Nikon Eclipse *Ti*) to form a magnetic tweezers platform, in conjunction with 4.5 μm diameter superparamagnetic beads. The microscope stage was immobilized, ensuring that the forces generated by the magnetic tweezers were identical in every image. Magnetic fields were generated by passing electrical current around copper coils wound around a mild steel core and focused over the sample using pole pieces on each side of the imaging region. Automated control of the field profile and direction was achieved with millisecond precision by powering the field from each pole piece independently, via a computer interface. Facing poles are separated by 36 mm, such that there is an 18 mm gap between the face of each pole and the centre of the imaging area.

The force acting on the superparamagnetic bead (which is transferred to the anchoring receptors) is determined by the magnetic properties of the beads and the spatial profile of the magnetic field (Bryan et al., 2010). To calibrate the magnetic field profile, a Gaussmeter (GM7, Hirst) was used to measure the field at each pole (at the centre of the surface facing the sample) and in the imaging position. More detailed calculations of the field profile were made by fitting these experimental data with a computational model generated with the ANSYS software package. Taking into account the three-dimensional nature of the electromagnet, the ANSYS program solved the Biot-Savart equation over a finite element mesh to calculate the field profile around the imaging region. Resultant forces from this field profile were calculated using the finite-element method described in (Bryan et al., 2010), such that the total force acting on the bead (*F*) at each position calculated is given by

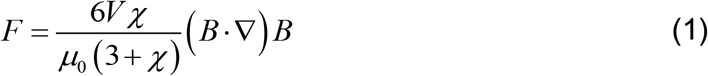

where V is the bead volume, *χ* (= 3.1) is the magnetic susceptibility of the bead, *μ*_0_ (= 4π×10^-7^ NA^-2^) is the permeability of free space, ∇ is the mathematical operator nabla, and *B* is the magnetic field at the calculated position.

For magnetic tweezers experiments, superparamagnetic beads were precipitated onto confluent HUVECs prepared on 1% fibronectin coated microdishes (μ-Dish 35 mm ibidi™) at a concentration of 250×10^4^ beads per 50×10^4^ cells per dish. Beads were incubated for 30 minutes and then unbound beads removed by exchange of media with sodium-buffered saline prior to the application of force. Bead movement was recorded using an inverted microscope (Nikon Eclipse *Ti*) coupled to a photometrics CoolSnap MYO camera and tracked using Spot Tracker plugin of ImageJ.

### Mice

ITGB1 was deleted from EC of adult mice (ITGB1 conditional knockout; ITGB1^cKO^). This was carried out by crossing mice containing a tamoxifen-inducible EC-specific Cre (Gothert et al., 2004; Mahmoud et al., 2016) (endothelial-SCL-Cre-ERT) with a strain containing a floxed version of *itgb1* (*itgb1^fl/fl^*). To activate Cre, tamoxifen (Sigma) was administered intraperitoneally for 5 consecutive days (100 mg/kg) (Mahmoud et al., 2016). ITGB1^cKO^ mice, aged 8-12 weeks, were culled 10 days after the first injection and systematically compared with control littermates treated under the same conditions. All mice were used in accordance with UK legislation (1986 Animals (Scientific Procedures) Act) and experiments were carried out under UK Home Office Project Licence (P28AA2253) for experimentation.

### *En face* staining of murine endothelium

The expression levels of specific proteins were quantified in EC at regions of the outer curvature (unidirectional flow; disease protected) or inner curvature (bidirectional flow; disease prone) of the murine aortic arch by *en face* staining. Animals were killed by I.P injection of pentobarbital. Aortae were perfused *in situ* with PBS and then perfusion-fixed with 4% paraformaldehyde prior to staining using specific primary antibodies and AlexaFluor568-conjugated secondary antibodies (Life Technologies) or with AlexaFluor568-phalloidin (ThermoFisher Scientific). EC were co-stained using anti-PECAM-1 antibody (clone: MEC 13.3, Biolegend), conjugated to FITC fluorophore. DAPI (Sigma) was used to identify nuclei. Stained vessels were analysed using super-resolution confocal microscopy (Zeiss LSM880 AiryScan Confocal). As experimental controls for specific staining, isotype matched monoclonal antibodies raised against irrelevant antigens were used. The expression of total and active β1 integrin was assessed by quantification of fluorescent intensity for multiple cells using ImageJ software (1.48v). Endothelial cell alignment was quantified by measuring the angle of the major axis of the nucleus in multiple cells (20 per field of view) as previously described (Poduri et al., 2017).

### Steered molecular dynamic (SMD) simulations

SMD simulations were performed with GROMACS 4.6 using the GROMOS96 53a6 force field. The αVβ3 ectodomain structure (PBD:3IJE) was used to run the simulations (Xiong et al., 2009). This structure was chosen because a 3D structure of the ectodomain of a β1-containing integrin is not available. Note that the αV 839 to αV 867 region that is missing from the crystal structure of the αVβ3 ectodomain is also missing from our model and that the unstructured regions αV 955 to αV 967 and β3 685 to β3 695 were removed for the simulations. The Parrinello-Rahman barostat (Parrinello and Rahman, 1981) was used for pressure coupling with isotropic pressure coupling. The V-rescale thermostat (Bussi et al., 2007) was used for temperature coupling. The Particle Mesh Ewald (PME) algorithm (Darden et al., 1993) was used to model long-range electrostatic interactions and the LINCS algorithm (Hess et al., 1997) was used to constrain bond lengths. The integrin was positioned in the simulation box with the Calf-1 and Calf-2 domains in a tilted (~30^°^) orientation relative to the xy plane (Fig. 3). This is believed to be the inactive orientation of an integrin. The simulation box size was ~29.4 x ~29.4 x ~19.3 nm and the system contained 1586961 atoms. The simulation system was solvated with waters and 150 nM of NaCl was added to neutralize the systems. Subsequently, the system was equilibrated for 2 ns with the protein Cα atoms restrained, followed by SMD simulations. A time step of 2 fs was used for the SMD simulations. The temperature of the simulations was 310 K. The GROMACS pull code was used (pull = constant-force) to apply a constant force on the center of mass of the βA domain. The force was parallel to the xy plane of the simulation box. During the SMD simulations the Cα atoms of the αV Calf-2 domain (residues:743-954) were restrained in all directions and the Cα atoms of the β3 β-tail domain (residues: 605-684) were restrained in the z direction to mimic the cell membrane. Three simulations were run using a force of 200 kJ mol^-1^ nm^-1^ for 80 ns each. Due to the size of the system ~1.5M particles, these forces, albeit higher to the force integrins may experience in the cell, enabled us to investigate the structural/molecular rearrangements that shear stress causes in the integrin at a reasonable timescale.

### Statistical analysis

Statistics were performed using paired Student’s ŕ-test or ANOVA (multiple comparisons) in GraphPad Prism 6. Differences between means were considered significant when p<0.05. Data are represented as means ± S.E.M. *p<0.05, **p<0.01, ***p<0.001.

## ACKNOWLEDGEMENTS

IX, CS, VR, JS-C, HR, SF, MTB and PCE were funded by the British Heart Foundation. MF was funded by Kidney Research UK. MJH was funded by Cancer Research UK (C13329/A21671). This work was undertaken on ARC2, part of the High Performance Computing facilities at the University of Leeds, UK. The authors declare no competing financial interests.

## AUTHOR CONTRIBUTIONS

P.C. Evans acquired funding, conceived and designed the study, supervised the work, analysed the data and wrote the manuscript. I. Xanthis, A. Kalli, V. Ridger, J. Waltho, M.J. Humphries and M.T. Bryan conceived and designed the study, acquired and analysed the data and edited the manuscript. C. Souilhol, J. Serbanovic-Canic, H. Roddie, M. Fragiadaki, R. Wong, D.R. Shah, J.A. Askari, L. Canham, N. Akhtar, S. Feng and E. Pinteaux made substantial contributions to interpreting the data and revising the manuscript for important intellectual content.

## REFERENCES

Ajami, N. E., Gupta, S., Maurya, M. R., Nguyen, P., Li, J. Y., Shyy, J. Y., Chen, Z., Chien, S. and Subramaniam, S. (2017). Systems biology analysis of longitudinal functional response of endothelial cells to shear stress. Proc Natl Acad Sci U S A 114, 10990–10995.

Albarran-Juarez, J., Iring, A., Wang, S., Joseph, S., Grimm, M., Strilic, B., Wettschureck, N., Althoff, T. F. and Offermanns, S. (2018). Piezo1 and Gq/G11 promote endothelial inflammation depending on flow pattern and integrin activation. J Exp Med 215, 2655–2672.

Ando J. and Yamamoto K. (2013). Flow detection and calcium signalling in vascular endothelial cells. Cardiovascular Research 99, 260–268.

Baeyens, N., Mulligan-Kehoe, M. J., Corti, F., Simon, D. D., Ross, T. D., Rhodes, J. M., Wang, T. Z., Mejean, C. O., Simons, M., Humphrey, J. et al. (2014). Syndecan 4 is required for endothelial alignment in flow and atheroprotective signaling. Proceedings of the National Academy of Sciences of the United States of America 111, 17308–17313.

Bhullar, I. S., Li, Y. S., Miao, H., Zandi, E., Kim, M., Shyy, J. Y. J. and Chien, S. (1998). Fluid shear stress activation of I kappa B kinase is integrin-dependent. Journal of Biological Chemistry 273, 30544–30549.

Bryan, M. T., Smith, K. H., Real, M. E., Bashir, M. A., Fry, P. W., Fischer, P., Im, M. Y., Schrefl, T., Allwood, D. A. and Haycock, J. W. (2010). Switchable Cell Trapping Using Superparamagnetic Beads. Ieee Magnetics Letters 1.

Buschmann, I., Pries, A., Styp-Rekowska, B., Hillmeister, P., Loufrani, L., Henrion, D., Shi, Y., Duelsner, A., Hoefer, I., Gatzke, N. et al. (2010). Pulsatile shear and Gja5 modulate arterial identity and remodeling events during flow-driven arteriogenesis. Development 137, 2187–2196.

Bussi, G., Donadio, D. and Parrinello, M. (2007). Canonical sampling through velocity rescaling. J Chem Phys 126, 014101.

Byron, A., Humphries, J. D., Askari, J. A., Craig, S. E., Mould, A. P. and Humphries, M. J. (2009). Anti-integrin monoclonal antibodies. J Cell Sci 122, 4009–11.

Carlson, T. R., Hu, H. Q., Braren, R., Kim, Y. H. and Wang, R. A. (2008). Cell-autonomous requirement for beta 1 integrin in endothelial cell adhesion, migration and survival during angiogenesis in mice. Development 135, 2193–2202.

Chen, J., Green, J., Yurdagul, A., Albert, P., McInnis, M. C. and Orr, A. W. (2015). alpha v beta 3 Integrins Mediate Flow-Induced NF-kappa B Activation, Proinflammatory Gene Expression, and Early Atherogenic Inflammation. American Journal of Pathology 185, 2575–2589.

Chen, K. D., Li, Y. S., Kim, M., Li, S., Yuan, S., Chien, S. and Shyy, J. Y. J. (1999). Mechanotransduction in response to shear stress - Roles of receptor tyrosine kinases, integrins, and Shc. Journal of Biological Chemistry 274, 18393–18400.

Chen, W., Lou, J. Z., Evans, E. A. and Zhu, C. (2012). Observing force-regulated conformational changes and ligand dissociation from a single integrin on cells. Journal of Cell Biology 199, 497–512.

Collins, C., Guilluy, C., Welch, C., O’Brien, E. T., Hahn, K., Superfine, R., Burridge, K. and Tzima, E. (2012). Localized Tensional Forces on PECAM-1 Elicit a Global Mechanotransduction Response via the Integrin-RhoA Pathway. Current Biology 22, 2087–2094.

Conforti, G., Dominguezjimenez, C., Zanetti, A., Gimbrone, M. A., Cremona, O., Marchisio, P. C. and Dejana, E. (1992). HUMAN ENDOTHELIAL-CELLS EXPRESS INTEGRIN RECEPTORS ON THE LUMINAL ASPECT OF THEIR MEMBRANE. Blood 80, 437–446.

Conforti, G., Zanetti, A., Jimenez, C. D., Cremona, O., Marchisio, P. C. and Dejana, E. (1991). HUMAN ENDOTHELIAL-CELLS EXPRESS AN INACTIVE ALPHA-V-BETA-3 RECEPTOR ON THEIR APICAL SURFACE WHOSE ACTIVITY CAN BE MODULATED. Thrombosis and Haemostasis 65, 962–962.

Darden, T., York, D. and Pedersen, L. (1993). PARTICLE MESH EWALD - AN N.LOG(N) METHOD FOR EWALD SUMS IN LARGE SYSTEMS. Journal of Chemical Physics 98, 10089–10092.

Feaver, R. E., Gelfand, B. D. and Blackman, B. R. (2013). Human haemodynamic frequency harmonics regulate the inflammatory phenotype of vascular endothelial cells. Nat Commun 4, 1525.

Ferraris, G. M., Schulte, C., Buttiglione, V., De Lorenzi, V., Piontini, A., Galluzzi, M., Podesta, A., Madsen, C. D. and Sidenius, N. (2014). The interaction between uPAR and vitronectin triggers ligand-independent adhesion signalling by integrins. Embo j 33, 2458–72.

Friedland, J. C., Lee, M. H. and Boettiger, D. (2009). Mechanically Activated Integrin Switch Controls alpha(5)beta(1) Function. Science 323, 642–644.

Givens C. and Tzima E. (2016). Endothelial Mechanosignaling: Does One Sensor Fit All? Antioxidants & Redox Signaling 25, 373–388.

Goel, R., Schrank, B. R., Arora, S., Boylan, B., Fleming, B., Miura, H., Newman, P. J., Molthen, R. C. and Newman, D. K. (2008). Site-specific effects of PECAM-1 on atherosclerosis in LDL receptor-deficient mice. Arterioscler Thromb Vasc Biol 28, 1996–2002.

Gothert, J. R., Gustin, S. E., van Eekelen, J. A. M., Schmidt, U., Hall, M. A., Jane, S. M., Green, A. R., Gottgens, B., Izon, D. J. and Begley, C. G. (2004). Genetically tagging endothelial cells in vivo: bone marrow-derived cells do not contribute to tumor endothelium. Blood 104, 1769–1777.

Harry, B. L., Sanders, J. M., Feaver, R. E., Lansey, M., Deem, T. L., Zarbock, A., Bruce, A. C., Pryor, A. W., Gelfand, B. D., Blackman, B. R. et al. (2008). Endothelial cell PECAM-1 promotes atherosclerotic lesions in areas of disturbed flow in ApoE-deficient mice. Arterioscler Thromb Vasc Biol 28, 2003–8.

Hess, B., Bekker, H., Berendsen, H. J. C. and Fraaije, J. (1997). LINCS: A linear constraint solver for molecular simulations. Journal of Computational Chemistry 18, 1463–1472.

Jalali, S., del Pozo, M. A., Chen, K., Miao, H., Li, Y., Schwartz, M. A., Shyy, J. Y. and Chien, S. (2001). Integrin-mediated mechanotransduction requires its dynamic interaction with specific extracellular matrix (ECM) ligands. Proc Natl Acad Sci U S A 98, 1042–6.

Kohler, R., Heyken, W. T., Heinau, P., Schubert, R., Si, H., Kacik, M., Busch, C., Grgic, I., Maier, T. and Hoyer, J. (2006). Evidence for a functional role of endothelial transient receptor potential V4 in shear stress-induced vasodilatation. Arterioscler Thromb Vasc Biol 26, 1495–502.

Kwak, B. R., Baeck, M., Bochaton-Piallat, M. L., Caligiuri, G., Daemens, M. J., Davies, P. F., Hoefer, I. E., Holvoet, P., Jo, H., Krams, R. et al. (2014). Biomechanical factors in atherosclerosis: mechanisms and clinical implications. European Heart Journal 35, 3013–+.

Lei, L., Liu, D. G., Huang, Y., Jovin, I., Shai, S. Y., Kyriakides, T., Ross, R. and Giordano, F. (2008). Endothelial Expression of Beta-1 Integrin Is Required for Embryonic Vascular Patterning and Postnatal Vascular Remodeling. Faseb Journal 22.

Li, J., Hou, B., Tumova, S., Muraki, K., Bruns, A., Ludlow, M. J., Sedo, A., Hyman, A. J., McKeown, L., Young, R. S. et al. (2014). Piezol integration of vascular architecture with physiological force. Nature 515, 279–U308.

Li, J., Su, Y., Xia, W., Qin, Y., Humphries, M. J., Vestweber, D., Cabanas, C., Lu, C. and Springer, T. A. (2017). Conformational equilibria and intrinsic affinities define integrin activation. Embo j 36, 629–645.

Li, S., Kim, M., Hu, Y. L., Jalali, S., Schlaepfer, D. D., Hunter, T., Chien, S. and Shyy, J. Y. (1997). Fluid shear stress activation of focal adhesion kinase. Linking to mitogen-activated protein kinases. J Biol Chem 272, 30455–62.

Loufrani, L., Retailleau, K., Bocquet, A., Dumont, O., Danker, K., Louis, H., Lacolley, P. and Henrion, D. (2008). Key role of alpha(1)beta(1)-integrin in the activation of PI3-kinase-Akt by flow (shear stress) in resistance arteries. Am J Physiol Heart Circ Physiol 294, H1906–13.

Luu, N. T., Glen, K. E., Egginton, S., Rainger, G. E. and Nash, G. B. (2013). Integrin-substrate interactions underlying shear-induced inhibition of the inflammatory response of endothelial cells. Thromb Haemost 109, 298–308.

Mahmoud, M. M., Kim, H. R., Xing, R. Y., Hsiao, S., Mammoto, A., Chen, J., Serbanovic-Canic, J., Feng, S., Bowden, N. P., Maguire, R. et al. (2016). TWIST 1 Integrates Endothelial Responses to Flow in Vascular Dysfunction and Atherosclerosis. Circulation Research 119, 450–+.

Matthews, B. D., Overby, D. R., Mannix, R. and Ingber, D. E. (2006). Cellular adaptation to mechanical stress: role of integrins, Rho, cytoskeletal tension and mechanosensitive ion channels. Journal of Cell Science 119, 508–518.

Matthews, B. D., Thodeti, C. K., Tytell, J. D., Mammoto, A., Overby, D. R. and Ingber, D. E. (2010). Ultra-rapid activation of TRPV4 ion channels by mechanical forces applied to cell surface beta1 integrins. Integr Biol (Camb) 2, 435–42.

Mendoza, S. A., Fang, J., Gutterman, D. D., Wilcox, D. A., Bubolz, A. H., Li, R., Suzuki, M. and Zhang, D. X. (2010). TRPV4-mediated endothelial Ca2+ influx and vasodilation in response to shear stress. Am J Physiol Heart Circ Physiol 298, H466–76.

Miyazaki, T., Honda, K. and Ohata, H. (2007). Requirement of Ca2+ influx- and phosphatidylinositol 3-kinase-mediated m-calpain activity for shear stress-induced endothelial cell polarity. Am J Physiol Cell Physiol 293, C1216–25.

Mould, A. P., Garratt, A. N., Askari, J. A., Akiyama, S. K. and Humphries, M. J. (1995). Identification of a novel anti-integrin monoclonal antibody that recognises a ligand-induced binding site epitope on the beta 1 subunit. FEBS Lett 363, 118–22.

Nguyen, B. A., V Suleiman, M. S., Anderson, J. R., Evans, P. C., Fiorentino, F., Reeves, B. C. and Angelini, G. D. (2014). Metabolic derangement and cardiac injury early after reperfusion following intermittent cross-clamp fibrillation in patients undergoing coronary artery bypass graft surgery using conventional or miniaturized cardiopulmonary bypass. Molecular and Cellular Biochemistry 395, 167–175.

Nordenfelt, P., Elliott, H. L. and Springer, T. A. (2016). Coordinated integrin activation by actin-dependent force during T-cell migration. Nat Commun 7, 13119.

Nordenfelt, P., Moore, T. I., Mehta, S. B., Kalappurakkal, J. M., Swaminathan, V., Koga, N., Lambert, T. J., Baker, D., Waters, J. C., Oldenbourg, R. et al. (2017). Direction of actin flow dictates integrin LFA-1 orientation during leukocyte migration. Nat Commun 8, 2047.

Orr, A. W., Ginsberg, M. H., Shattil, S. J., Deckmyn, H. and Schwartz, M. A. (2006). Matrix-specific suppression of integrin activation in shear stress signaling. Molecular Biology of the Cell 17, 4686–4697.

Orr, A. W., Sanders, J. M., Bevard, M., Coleman, E., Sarembock, I. J. and Schwartz, M. A. (2005). The subendothelial extracellular matrix modulates NF-kappa B activation by flow: a potential role in atherosclerosis. Journal of Cell Biology 169, 191–202.

Parrinello M. and Rahman A. (1981). POLYMORPHIC TRANSITIONS IN SINGLECRYSTALS - A NEW MOLECULAR-DYNAMICS METHOD. Journal of Applied Physics 52, 7182–7190.

Petridou N. I. and Skourides P. A. (2016). A ligand-independent integrin beta1 mechanosensory complex guides spindle orientation. Nat Commun 7, 10899.

Poduri, A., Chang, A. H., Raftrey, B., Rhee, S., Van, M. and Red-Horse, K. (2017). Endothelial cells respond to the direction of mechanical stimuli through SMAD signaling to regulate coronary artery size. Development 144, 3241–3252.

Puklin-Faucher, E., Gao, M., Schulten, K. and Vogel, V. (2006). How the headpiece hinge angle is opened: new insights into the dynamics of integrin activation. Journal of Cell Biology 175, 349–360.

Puklin-Faucher E. and Sheetz M. P. (2009). The mechanical integrin cycle. Journal of Cell Science 122, 179–186.

Shyy J. Y. and Chien S. (2002). Role of integrins in endothelial mechanosensing of shear stress. Circulation Research 91, 769–775.

Sorescu, G. P., Song, H., Tressel, S. L., Hwang, J., Dikalov, S., Smith, D. A., Boyd, N. L., Platt, M. O., Lassegue, B., Griendling, K. K. et al. (2004). Bone morphogenic protein 4 produced in endothelial cells by oscillatory shear stress induces monocyte adhesion by stimulating reactive oxygen species production from a nox1-based NADPH oxidase. Circulation Research 95, 773–779.

Su, Y., Xia, W., Li, J., Walz, T., Humphries, M. J., Vestweber, D., Cabanas, C., Lu, C. and Springer, T. A. (2016). Relating conformation to function in integrin alpha5beta1. Proc Natl Acad Sci U S A 113, E3872–81.

Suo, J., Ferrara, D. E., Sorescu, D., Guldberg, R. E., Taylor, W. R. and Giddens, D. P. (2007). Hemodynamic shear stresses in mouse aortas: implications for atherogenesis. Arteriosclerosis, Thrombosis, and Vascular Biology 27, 346–51.

Tanjore, H., Zeisberg, E. M., Gerami-Naini, B. and Kalluri, R. (2008). beta 1 integrin expression on endothelial cells is required for angiogenesis but not for vasculogenesis. Developmental Dynamics 237, 75–82.

Thodeti, C. K., Matthews, B., Ravi, A., Mammoto, A., Ghosh, K., Bracha, A. L. and Ingber, D. E. (2009). TRPV4 Channels Mediate Cyclic Strain-Induced Endothelial Cell Reorientation Through Integrin-to-Integrin Signaling. Circulation Research 104, 1123–U278.

Tzima, E., del Pozo, M. A., Shattil, S. J., Chien, S. and Schwartz, M. A. (2001). Activation of integrins in endothelial cells by fluid shear stress mediates Rho-dependent cytoskeletal alignment. Embo Journal 20, 4639–4647.

Tzima, E., Irani-Tehrani, M., Kiosses, W. B., Dejana, E., Schultz, D. A., Engelhardt, B., Cao, G., DeLisser, H. and Schwartz, M. A. (2005). A mechanosensory complex that mediates the endothelial cell response to fluid shear stress. Nature 437, 426–31.

Urbich, C., Dernbach, E., Reissner, A., Vasa, M., Zeiher, A. M. and Dimmeler, S. (2002). Shear stress-induced endothelial cell migration involves integrin signaling via the fibronectin receptor subunits alpha(5) and beta(1). Arterioscler Thromb Vasc Biol 22, 69–75.

Urbich, C., Fritzenwanger, M., Zeiher, A. M. and Dimmeler, S. (2000). Laminar shear stress upregulates the complement-inhibitory protein clusterin - A novel potent defense mechanism against complement-induced endothelial cell activation. Circulation 101, 352–355.

Wang, C., Baker, B. M., Chen, C. S. and Schwartz, M. A. (2013). Endothelial cell sensing of flow direction. Arterioscler Thromb Vasc Biol 33, 2130–6.

Wang, N. P., Miao, H., Li, Y. S., Zhang, P., Haga, J. H., Hu, Y. L., Young, A., Yuan, S. L., Nguyen, P., Wu, C. C. et al. (2006). Shear stress regulation of Kruppel-like factor 2 expression is flow pattern-specific. Biochemical and Biophysical Research Communications 341, 1244–1251.

Warboys, C. M., de Luca, A., Amini, N., Luong, L., Duckles, H., Hsiao, S., White, A., Biswas, S., Khamis, R., Chong, C. K. et al. (2014). Disturbed Flow Promotes Endothelial Senescence via a p53-Dependent Pathway. Arteriosclerosis Thrombosis and Vascular Biology 34, 985–995.

Wu, W., Xiao, H., Laguna-Fernandez, A., Villarreal, G., Wang, K. C., Geary, G. G., Zhang, Y. Z., Wang, W. C., Huang, H. D., Zhou, J. et al. (2011). Flow-Dependent Regulation of Kruppel-Like Factor 2 Is Mediated by MicroRNA-92a. Circulation 124, 633–U231.

Xiong, J. P., Mahalingham, B., Alonso, J. L., Borrelli, L. A., Rui, X., Anand, S., Hyman, B. T., Rysiok, T., Muller-Pompalla, D., Goodman, S. L. et al. (2009). Crystal structure of the complete integrin alphaVbeta3 ectodomain plus an alpha/beta transmembrane fragment. J Cell Biol 186, 589–600.

Yang, B. H., Radel, C., Hughes, D., Kelemen, S. and Rizzo, V. (2011). p190 RhoGTPase-Activating Protein Links the beta 1 Integrin/Caveolin-1 Mechanosignaling Complex to RhoA and Actin Remodeling. Arteriosclerosis Thrombosis and Vascular Biology 31, 376–U300.

Zhu, J., Luo, B. H., Xiao, T., Zhang, C., Nishida, N. and Springer, T. A. (2008). Structure of a complete integrin ectodomain in a physiologic resting state and activation and deactivation by applied forces. Mol Cell 32, 849–61.

Zovein, A. C., Luque, A., Turlo, K. A., Hofmann, J. J., Yee, K. M., Becker, M. S., Fassler, R., Mellman, I., Lane, T. F. and Iruela-Arispe, M. L. (2010). beta 1 Integrin Establishes Endothelial Cell Polarity and Arteriolar Lumen Formation via a Par3-Dependent Mechanism. Developmental Cell 18, 39–51.

